# Evolutionary constraint genes implicated in autism spectrum disorder across 2,054 nonhuman primate genomes

**DOI:** 10.1101/2023.11.08.566198

**Authors:** Yukiko Kikuchi, Mohammed Uddin, Joris A. Veltman, Sara Wells, Marc Woodbury-Smith

## Abstract

**Background:** Significant progress has been made in elucidating the genetic underpinning of Autism Spectrum Disorder (ASD). This childhood-onset chronic disorder of cognition, communication and behaviour ranks among the most severe from a public health perspective, and it is therefore hoped that new discoveries will lead to better therapeutic options. However, there are still significant gaps in our understanding of the link between genomics, neurobiology and clinical phenotype in scientific discovery. New models are therefore needed to address these gaps. Rhesus macaques (*Macaca mulatta*) have been extensively used for preclinical neurobiological research because of remarkable similarities to humans across biology and behaviour that cannot be captured by other experimental animals.

**Methods:** We used the macaque Genotype and Phenotype (mGAP) resource (v2.0) consisting of 2,054 macaque genomes to examine patterns of evolutionary constraint in known human neurodevelopmental genes. Residual variation intolerance scores (RVIS) were calculated for all annotated autosomal genes (N = 18,168) and Gene Set Enrichment Analysis (GSEA) was used to examine patterns of constraint across ASD genes and related neurodevelopmental genes.

**Results:** We demonstrated that patterns of constraint across autosomal genes are correlated in humans and macaques, and that ASD-implicated genes exhibit significant constraint in macaques (*p* = 9.4 x 10^-27^). Among macaques, many key ASD genes were observed to harbour predicted damaging mutations. A small number of key ASD genes that are highly intolerant to mutation in humans, however, showed no evidence of similar intolerance in macaques (*CACNA1D*, *CNTNAP2*, *MBD5*, *AUTS2* and *NRXN1*). Constraint was also observed across genes implicated in intellectual disability (*p* = 1.1 x 10^-46^), epilepsy (*p* = 2.1 x 10^-33^) and schizophrenia (*p* = 4.2 x 10^-45^), and for an overlapping neurodevelopmental gene set (*p* = 4.0 x 10^-10^)

**Limitations:** The lack of behavioural phenotypes among the macaques whose genotypes were studied means that we are unable to further investigate whether genetic variants have similar phenotypic consequences among nonhuman primates.

**Conclusion:** The presence of pathological mutations in ASD genes among macaques, and the evidence of similar constraints in these genes to humans, provide a strong rationale for further investigation of genotype-phenotype relationships in nonhuman primates. This highlights the importance of identifying phenotypic behaviours associated with clinical symptoms, elucidating the neurobiological underpinnings of ASD, and developing primate models for translational research to advance approaches for precision medicine and therapeutic interventions.

## Introduction

Autism Spectrum Disorder (ASD) is a clinically defined, early-onset lifelong neurodevelopmental disorder that affects social, cognitive and communicative abilities. The health and well-being of autistic people is a priority among many governments worldwide (e.g., in the UK: Think Autism, 2014; The NHS Long Term Plan, 2019). However, despite recent advances in ASD genetics, there are currently no effective biologically-derived treatment options that improve the health and wellbeing of autistic people. This underscores the importance of developing new animal models that can facilitate an understanding of both genetic and biological factors, which will inform and advance the efficacy and precision of potential therapeutic options to improve clinical outcomes.

Rhesus macaques (*Macaca mulatta*) have been extensively used for preclinical neurobiological research and macaques and humans share remarkable similarities in social and cognitive behaviours, and neurobiology that cannot be captured by other experimental animals (1–4). More recently, primate species have been used to infer deleteriousness of mutations in humans, thereby facilitating better diagnostic accuracy in patients with genetic diseases (5). Rodent models, although powerful, are insufficient as they lack primate-specific frontal structure that is critical for social and cognitive behaviours associated with ASD. Consequently, a primate model could provide important information to clarify the evolutionary divergent mechanisms that occurred between rodent models and humans. For instance, macaques have forward-looking eyes and their eye movement patterns are similar to those of typically developing children who attend preferentially to features of social or emotionally arousing stimuli (6–9); such gaze behaviours are abnormal in ASD (10–13). Furthermore, social behaviours among macaques can be quantitatively assessed and characterised in a way similar to the screening for ASD in humans, by using assessments such as the macaque Social Responsiveness Scale (SRS)-Revised (mSRS-R) (14,15), which is derived from the human SRS (16,17). Consistent with the distribution of SRS scores in the general human population, the scores among macaques are also continuously distributed. Moreover animals with SRS scores greater than 1.5 SD above the mean exhibit a greater burden of autistic-like traits (15).

Genetically altered nonhuman primate models of ASD have been generated to target single-genes that are implicated in ASD, such as *SHANK3* (18,19) and *MECP2* (20,21) in which abnormal social and repetitive behaviours, sleep disturbance and impairments in cognitive function are observed. Although single gene knockout models may work well for characterising the pathological mechanisms in individual genes, given the polygenic nature of ASD, genome-wide studies are crucial. It is advantageous that there is now a robust rhesus reference genome which provides a framework for evaluating mutational characteristics among macaque genes that are implicated in human pathology. Recently a large macaque genomic database, the macaque Genotype and Phenotype (mGAP) resource, has become available, providing a catalogue of annotated macaque variant data derived from exome and whole genome sequencing (22).

In this study, using the mGAP resource, we examined genetic constraint among ASD-implicated autosomal genes across 2,054 macaque genomes. We hypothesized that macaques would show similar genetic constraint to humans among orthologous genes, consistent with their likely importance in brain development, and consequently consistent with the likelihood of pathological consequences from those variants predicted to be damaging. As a corollary, we extended our analyses to examine constraint among other neurodevelopmental genes implicated in epilepsy, intellectual disability (ID) and schizophrenia.

## Methods

### Rhesus macaque variant catalog (mGAP)

The Macaque Genotype and Phenotype Resource (mGAP) (https://mgap.ohsu.edu/) is an open access database of exome and whole genome sequencing from macaques housed at a number of primate research centres, but principally from a multi-generational outbred cohort of macaques bred at the Oregon National Primate Research Center. The iteration used in our analyses (v2.0) includes data from 2,054 Indian and Chinese-origin macaques, with additional limited phenotypes comprising basic metabolic parameters. Sequence data are analysed by the mGAP team using a modified version of the Broad Institute/GATK SNP and Indel Discovery Best Practices adapted for use with macaque data. All v2.0 data are aligned to the Mmul_10 reference genome. The mGAP v2.0 release contains variants passing a quality filtering process including the use of Mendelian inheritance for validation.

### Clinical gene sets

To derive clinical gene sets for ASD and other NNDs (i.e., epilepsy, intellectual disability [ID], and schizophrenia [SZ]), we used the DisGeNET database of curated genes and variants (https://www.disgenet.org/). These genes and variants are collated from different data sources, including those extracted by text mining the scientific literature (23). Genes and variants are annotated in various ways, including the addition of metrics that demonstrate the strength of disease-gene/variant associations. DisGeNET is regularly updated (October 2021 update used for current analyses) and provides metrics that allow researchers to classify genes according to level of evidence. The final lists of DisGeNET genes comprised 793 ASD genes, 1,570 ID genes, 871 epilepsy genes and 2,018 schizophrenia genes. Separately, we used the latest version of the Simons Foundation Autism Research Initiative (SFARI) SFARI-Gene platform to download a list of ASD-implicated genes (24). This list comprised 729 ASD genes. There were 292 genes occurring in both SFARI and DisGeNET ASD gene lists.

### Quality screening of SNVs

We used bcftools, vcftools and R scripts for all analyses (all R scripts are available in our GitHub repository, and related files are available as Supplementary Files). We first generated summary statistics for variant quality and variant depth and excluded all variants with a quality score of < 30, and all variants whose depth was < 10 or > 50. We did not exclude variants by their minor allele frequency (MAF) to avoid excluding any variants that might potentially be of significance. For the purpose of this current analysis, we focussed on only single nucleotide variants (SNVs), therefore excluding indels from all downstream analyses. We also focussed on only the autosomes.

### Genetic constraint in human and macaque genome

#### Residual Variation Intolerance Score (RVIS) analysis

To investigate whether genes implicated in ASD and other NDDs were subject to constraint against the accumulation of damaging mutations in the macaque genome, we generated gene-by-gene Residual Variation Intolerance Scores (RVIS) for macaque genomes, which uses an intolerance ranking system for each gene as described in Petrovski et al. (2013) (25). Briefly, this constraint metric is calculated by regressing the total number of functional variants (i.e., common missense and loss of function SNVs) in a gene on the total number of protein-coding variants in that gene. The studentized residuals (mean = 0) are then taken as the RVIS, which provides a measure that captures departure from the average mutational burden. When RVIS = 0, the gene has the average number of common variants given its total mutational burden; when RVIS < 0, the gene has fewer common variations than predicted (i.e., more constrained, mutation intolerant); when RVIS > 0, it has more common variation than predicted (i.e., less constrained, mutation tolerant). Among macaque data, two genes (*MUC16*, *LOC106997451*) with RVIS scores greater than 11 were removed from the analysis as potential outliers. Neither of these genes is deemed ‘essential’ according to Blomen et al.’s (26) and Godini at al.’s (27) ‘essentialome’, and neither feature in the SFARI or DisGeNET gene lists for the neurodevelopmental disorders investigated (see below). Applying the same criterion to human RVIS data (25) resulted in 5 genes (*MUC16*, *MUC17*, *MUC5B*, *AHNAK2*, *FLG*) being excluded.

We undertook analysis of group differences in RVIS score using t-tests. In order to examine the magnitude of constraint differences between groups, we also undertook logistic regression. All analyses were implemented in R, with the code available in our repository.

#### Gene Set Enrichment Analysis (GSEA) analysis

We next undertook Gene Set Enrichment Analysis (GSEA) using the hypeR package in R (28). hypeR implements the Kolmogorov-Smirnov test to determine whether a set of genes (in our case, our curated ASD and NDD genes) is randomly distributed across a ranked list (in our case, all autosomal genes), or instead, is preferentially clustered towards either end, as measured by calculation of an ‘enrichment score (ES)’. Through permutation of the ranked data, a null distribution of enrichment scores can be generated allowing a nominal p-value to be produced. As a corollary, we also undertook a gene set over-representation analysis using the same hypeR package. This method implements a hypergeometric test, which we used to examine the over-representation of ASD and other NDD gene sets among the top 2% constrained genes. These constrained genes were identified by first ranking genes and then identifying the top 363 genes (18,166 * 0.02).

### Analysis of predicted-damaging mutations

Where possible, variants in mGAP have been lifted to the human genome (hg38) and annotated using a variety of bioinformatic tools, as described on the mGAP website. In this way, a list of predicted damaging variants has been generated by the mGAP team, comprising those variants that either (i) are identical to a ClinVar allele annotated as pathogenic, (ii) are predicated damaging according to Polyphen2 score, or (iii) predicted as high impact by SnpEff. Using the mGAP ‘predicted_damaging’ file for v2.0, we examined both (i) the number of ASD-implicated genes (SFARI and DisGeNET curated) that harboured one or more predicted-damaging mutations in macaques and (ii) whether any of these predicted damaging mutations were identical to ASD mutations listed in DisGeNET and SFARI. The UCSC’s LiftOver tool was used to change coordinates to and from the macaque MMul_10 and hg38 builds. Scripts used for these analyses are all available.

## Results

We undertook analyses on mGAP release 2.0, made available on the 18^th^ January 2021. All samples are aligned to the Mmul_10 reference genome, and variants called using GATK’s GenomicsDB pipeline as detailed on the mGAP website. This release included 2,054 macaques, among whom basic phenotype data are available on 1,299 indicating the majority to be of Indian origin (N = 1,206) with the remainder Chinese (N = 13), mixed Indian/Chinese (N = 8) or unknown origin (N = 71). There were 703 male and 596 female macaques among those with available sex data. We first filtered variants according to variant quality and depth as described in the Methods. After filtering, a total of 37,503,663 variants remained across the 20 macaque autosomes comprising 32,507,418 SNPs (86.7 %) and 4,996,245 Indels (13.3 %). The total number of variants annotated missense according to SnpEff is 370,683. **Supplementary Table 1** shows metrics before and after our filtering steps. We excluded Indels and undertook analysis only on SNPs.

Genetic constraint was calculated using RVIS across all 18,168 genes as described in Methods. Using human RVIS data from Petrovski et al., (2013) (25), we compared the RVIS in 18,166 genes in the human and macaque genome and identified a significant correlation between human and macaque RVIS scores (**Supplementary Figure 1**, Pearson’s product-moment correlation = 0.39 [95% CI = 0.38 - 0.41], *P* < 2.2e^-16^). To ensure the validity of RVIS as a measure of constraint, we also investigated whether the RVIS scores in macaques correctly identify those genes that are deemed ‘essential’ in humans, as identified by Blomen et al. (1,734 genes) (26) and Godini et al. (27) (1,586 genes) (**Supplementary File 1**: Blomen-2015-core-essentialome; **Supplementary File 2**: Godini -2015-essential-genes). Using logistic regression, we identified that in both instances the essential genes were significantly more constrained than non-essential genes (‘Blomen genes’: Beta = -0.17, SE = 0.04, *P* = 1.38e^-05^; ‘Godini genes’: Beta = -0.26, SE = 0.04, *P* = 4.19e^-11^).

### Genetic constraint of ASD implicated genes

We examined the distribution of RVIS scores among SFARI (**Figures 1A, 1C**) and DisGeNET (**Figures 1B, 1D**) ASD genes respectively. In both instances, the RVIS scores were indicative of greater constraint among ASD genes compared to the background genome (SFARI: N = 729, mean = -0.44, SD = 1.10; non-ASD: N = 17,437, mean = 0.02, SD = 0.67, *P* = 9.4 x 10^-27^, OR = 0.53; DisGenet: N = 793, mean = -0.32, SD = 0.88; non-ASD: N = 17,373, mean = 0.01, SD = 0.67, *P* = 4.6 x 10^-24^, OR = 0.61). (**Supplementary Table 2**)

**Figure 1.**
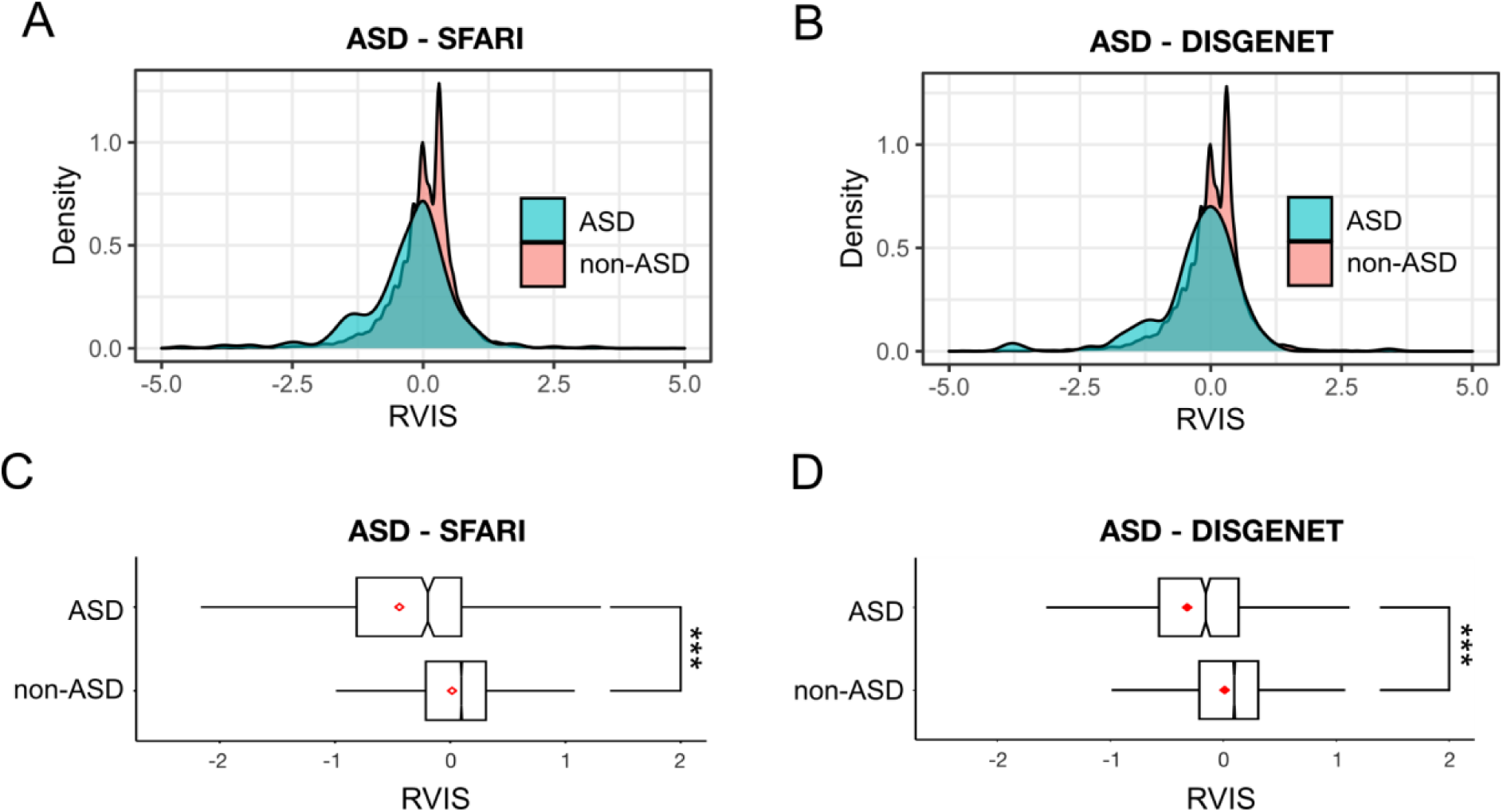
**A**. Density histogram of the Residual Variation Intolerance Scores (RVIS) for SFARI ASD genes (blue, N = 729) and non-ASD (red, N = 17,437) genes. The RVIS scores are normalised and hence the centre on a mean of 0. Delimiters on x-axis range from -5.0 to 5.0. **B**. The descriptions are the same as **A** for the DISGENET ASD genes (blue, N = 793) and non-ASD (red, N = 17,373) genes. **C**. Box and whisker plots depicting the distribution of RVIS scores with median (red line) for SFARI ASD and non-ASD genes. Delimiters on x-axis range from -1.5 to 1.5. *** *P* & 0.001 (t test). **D**. Same as C using the DisGenet ASD genes.

Next, Gene Set Enrichment Analysis (GSEA) was performed to investigate whether ASD genes were enriched among the most constrained genes ranked according to their RVIS. As expected, ASD genes showed evidence of enrichment among the more constrained genes (ES for ASD-SFARI = 0.23; ES for ASD-DisGenet = 0.19, **Figure 2 A-B**). As a corollary, we also investigated whether ASD genes were over-represented among the top 2% constrained genes (i.e., those genes with most negative RVIS, N = 369 genes) (**Figure 2C**). In each case, significant over-representation was observed in the ASD genes (ASD-DisGeNet, *P* < 9.6 x 10^-13^, ASD-SFARI, *P* < 2.8 x 10^-24^, **Supplementary Table 3**).

**Figure 2.**
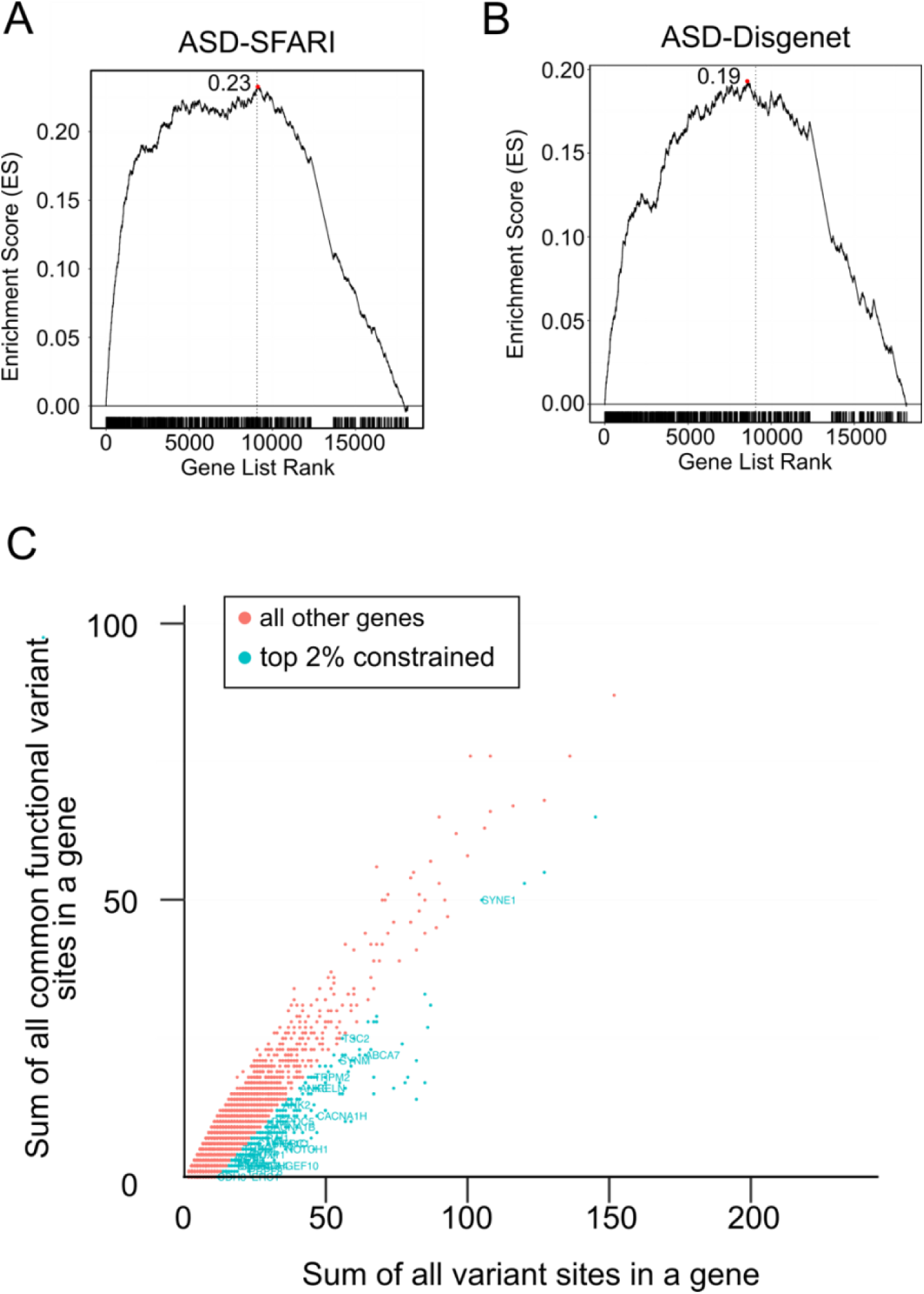
Gene Set Enrichment Analysis (GSEA) using ASD SFARI gene set (**A**) and DisGeNET ASD gene set (**B**). The x-axis denotes all genes ranked by the RVIS. The black vertical line indicates the ASD set genes. The y-axis is the Enrichment Score (ES) which represents the degree (i.e., the maximum deviation from zero) to which a set of genes is over-represented at the top of the ranked list accordingly to the RVIS. **C.** The scatter plot of RVIS scores for ASD-implicated genes. The blue plots are for the 2% extreme of the distribution i.e., most intolerant in which the annotated plot indicates the DisGeNET ASD set genes. The red plots indicate 98 % of the rest of the genes.

Next, we investigated whether any of the predicted damaging mutations identified by mGAP’s annotation pipeline (N = 128,705 variants) overlapped with known ASD variants as curated by DisGeNET (N=332 variants). The result shows that none of the macaque predicted damaging variants overlapped the known ASD variants. However, the majority of ASD genes included at least one predicted-damaging variant (**Table 1**).

**Table 1.** Counts of predicted damaging variants that overlap variants implicated for each studied NDD.

|  | Number of genes in DisGeNeT database | Number with one or more mGAP predicted damaging variants | Proportion of disease gene with one or more predicted damaging variant |
| --- | --- | --- | --- |
| ASD | 1,071 | 823 | 76.8% |
| ID | 2,165 | 1,732 | 80.0% |
| Epilepsy | 1,215 | 933 | 76.8% |
| Schizophrenia | 2,872 | 2,098 | 73.1% |

### Genetic constraint among other neurodevelopmental disorders

Next, we examined constraint scores for other NDD gene sets to test whether RVISs for different NDD gene sets were consistent with the evidence of greater constraint. The RVIS scores were as follows: schizophrenia (mean: -0.25, SD: 0.85), ID (mean: -0.30, SD: 0.84) and epilepsy (mean: -0.33, SD: 0.79). These differences were significant for all disorders studied (**Supplementary Table 2**). The RVISs were indicative of greater constraint across all gene sets (ID: p = 1.1 x 10^-46^, OR = 0.58; epilepsy: p = 2.1 x 10^-33^ OR = 0.60; schizophrenia: p = 4.2 x 10^-45^, OR = 0.61, **Figure 3**).

**Figure 3.**
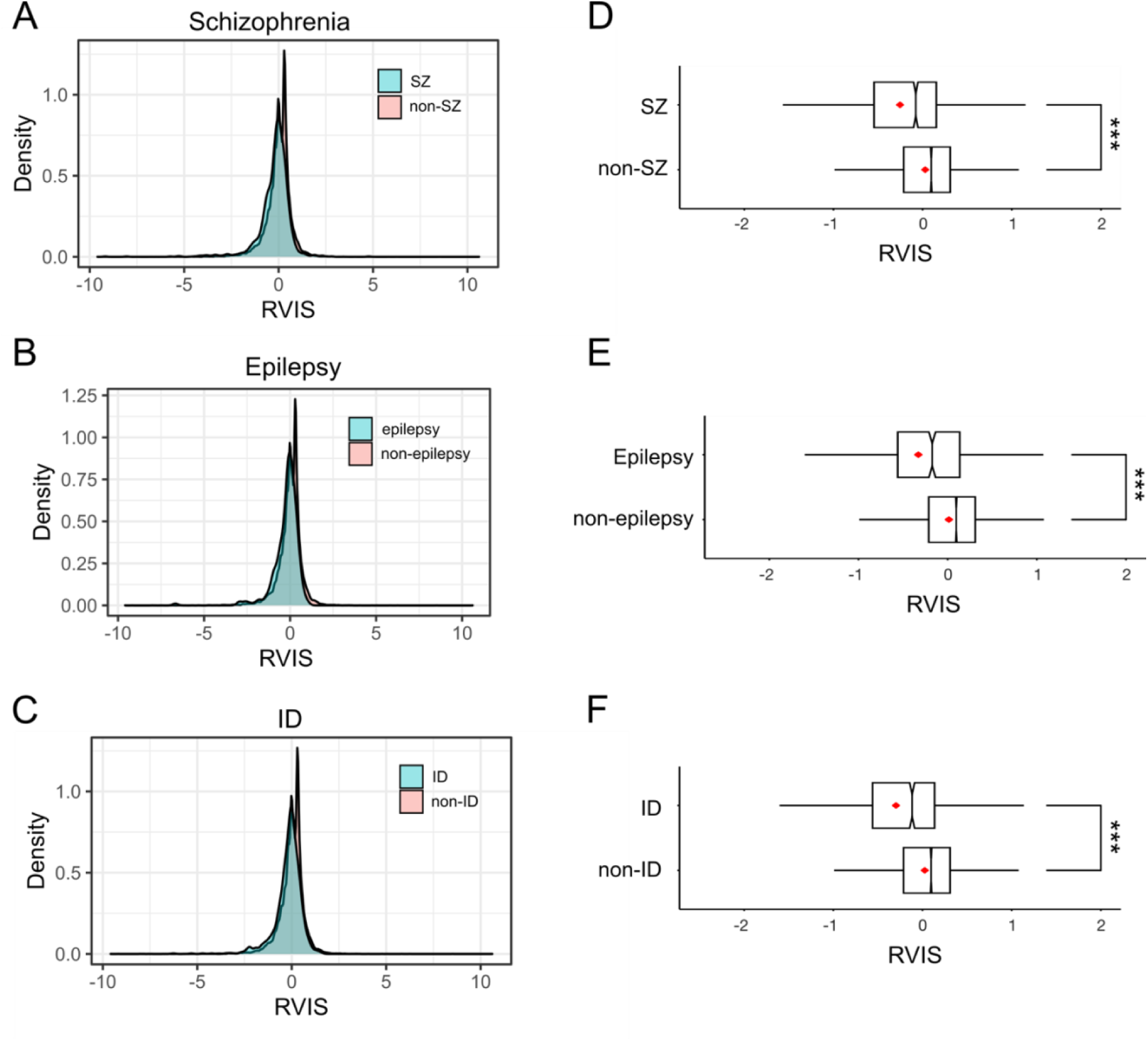
**A**. Density histogram of the Residual Variation Intolerance Scores (RVIS) for DisGeNET schizophrenia (**A**), epilepsy (**B**) and intellectual disability (ID) genes (**C**) (coloured in blue) and non-disorder implicated genes (red). The RVIS scores are normalised and hence the centre on a mean of 0. Delimiters on x-axis range from -5.0 to 5.0. **B**. The descriptions are the same as **A**. **D-F**. Box and whisker plots depicting the distribution of RVIS scores with median (red line) for schizophrenia (**D**), epilepsy (**E**) and ID (**F**) DisGeNET genes. Delimiters on x-axis range from -1.5 to 1.5. *** *P* & 0.001 (*t*-test).

We also undertook GSEA on these neurodevelopmental disorders. Consistent with our prediction, all gene sets showed evidence of enrichment towards the more constrained ‘end’ of the ranked genes (ES: 0.17 for schizophrenia; 0.2 for epilepsy; 0.19 for ID, **Figure 4**, **Supplementary Table 4** for each diagnosis). As a corollary, we also investigated whether NDD genes for each diagnosis were over-represented among the top 2% constrained genes. In each case, significant over-representation was observed (ID: *P* < 1.7 x 10^-19^, Epilepsy: *P* < 8.9 x 10^-12^, Schizophrenia: *P* < 5.9 x 10^-16^; **Supplementary Table 3**).

**Figure 4.**
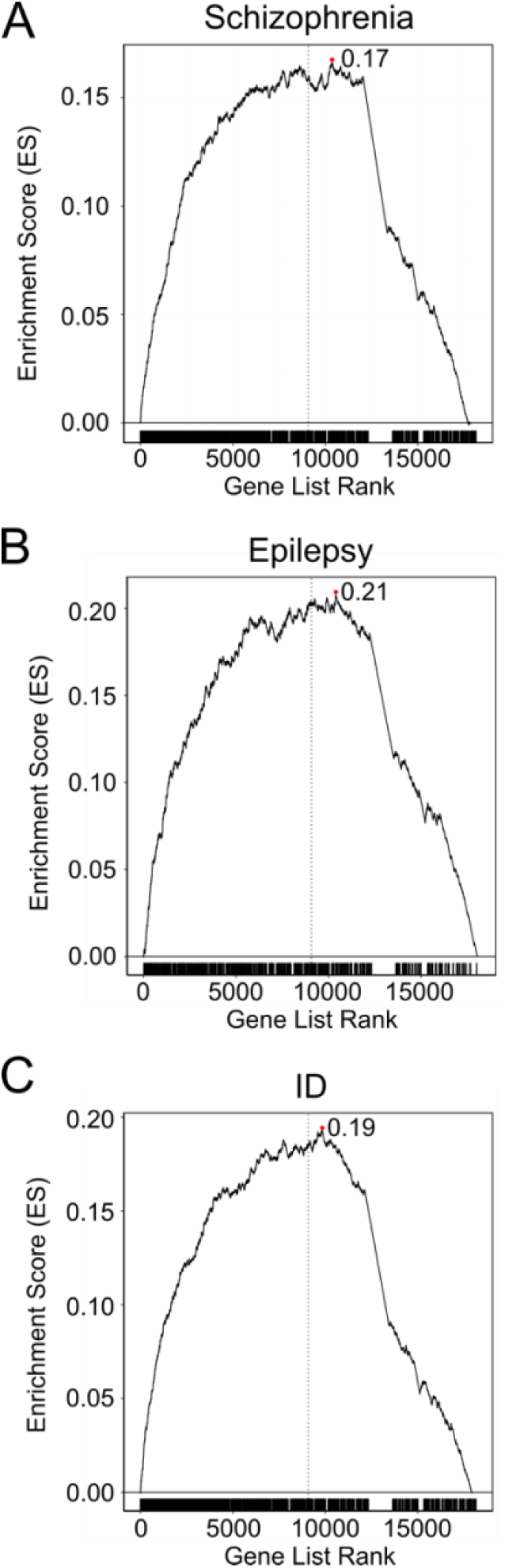
**A**. GSEA results using the set genes for schizophrenia (A), epilepsy (B) and ID (C). The descriptions are the same as Figure 2A.

Next, to investigate whether any of the predicted damaging mutations identified by mGAP’s annotation pipeline (N = 128,705) overlapped with known NDD variants, as curated by DisGeNET, we examined the frequency of the predicted damaging mutations. After lifting over co-ordinates, two predicted damaging variants are known schizophrenia variants in humans (rs2241621: 81270732 [hg38], *STON2*, A>C, T, missense variant; rs16969968: 78590583 [hg38], *CHRNA5*, G>A, missense variant), one is a known ID variant (rs587777623: 686986 [hg38], *DEAF1*, G>A, missense variant) and one a known epilepsy variant (rs28941773: 120739168 [hg38], *ACADS*, C>T, missense variant). As was the case for ASD genes (see above), the majority of genes across these disorders harboured one or more predicted-damaging mutation (ID: 80.0 %, epilepsy: 76.8 %, schizophrenia: 73.1%, **Table 1**).

### Genetic constraint among key NDD genes

Given the inclusive nature of these NDD gene sets, i.e., the relatively low threshold for any particular gene to be identified as disorder-implicated, we further refined our gene list to comprise only those genes that overlapped among all four disorders (N=101 genes, see **Supplementary File 3** – Overlapping NDD genes). The importance of the genes in this list is demonstrated by their key role in certain brain processes (**Supplementary File 4** - g:profiler mmulatta_intersections.csv).

**Figure 5** shows the distribution of RVIS scores in NDD-implicated genes and non-NDD genes (NDD: N = 101, mean = -0.52, SD: 0.75; non-NDD: N = 18,065, mean = 0.00, SD: 0.68, *P* = 4.0 x 10^-10^, OR = 0.59). (Supplementary Table 2)

**Figure 5.**
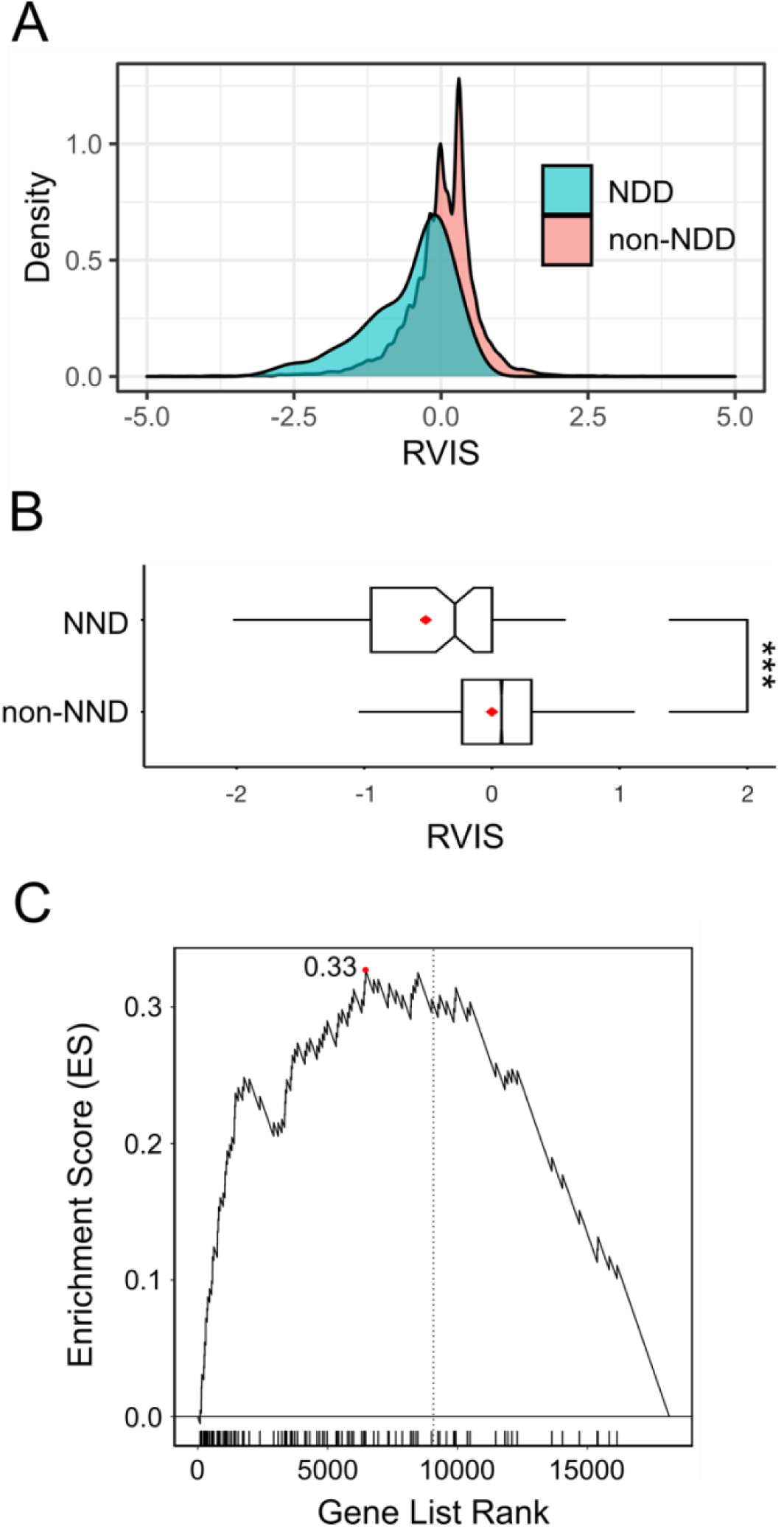
**A.** Density histogram of the Residual Variation Intolerance Scores (RVIS) for NDD (N = 101, blue) and non-NDD (N = 18,065, red) genes. The RVIS scores are normalised and hence the centre on a mean of 0. **B.** The box plots for any NDD genes (i.e., ASD, epilepsy, ID, schizophrenia) and non-NDD genes. The descriptions are the same as Figure 2C and D. Box plots **C.** GSEA for the overlapped NDD genes.

We examined the burden of predicted damaging mutations in these genes in the macaque data. In total, 91 of the 101 genes (90.1 %) had one or more predicted damaging variants, with a total of 809 variants across all genes (**Supplementary File 5** – Overlap Hits). We conclude, therefore, that despite the constraint against mutation among these NDD-implicated genes, macaques still harbour predicted damaging mutations, with potential phenotypic consequences. By way of illustration, **Table 2** shows the 20 predicted damaging variants with the largest Combined Annotation Dependent Depletion (CADD) scores (29). These scores are based on the integration of multiple annotation metrics into one score which earmarks likely pathogenicity. A score of 30 or more are deemed likely deleterious, indexing the most 0.1% pathogenic substitutions across the genome. Similarly, a score of 20 or more indexes the top 1% deleterious substitutions. Given these high CADD scores, and the fact that these genes are largely well-recognised as key NDD genes, it might be predicted that in humans these mutations would have clinical neurodevelopmental consequences.

**Table 2.** Predicted damaging variants in NDD genes and their CADD scores.

| Gene | Chr | Start (hg19) | End (hg19) | CADD |
| --- | --- | --- | --- | --- |
| *RAI1* | chr16 | 17625726 | 17625726 | 45 |
| *CNTN2* | chr1 | 62585295 | 62585295 | 40 |
| *MAPK1* | chr10 | 31173015 | 31173015 | 37 |
| *CACNA1D* | chr2 | 109940657 | 109940657 | 37 |
| *SLC52A2* | chr8 | 144641018 | 144641018 | 37 |
| *LRP2* | chr12 | 56308216 | 56308216 | 36 |
| *PIK3CG* | chr3 | 133216881 | 133216881 | 36 |
| *CNTN2* | chr1 | 62590334 | 62590334 | 35 |
| *ERBB4* | chr12 | 98730326 | 98730326 | 35 |
| *GRIK4* | chr14 | 113885581 | 113885581 | 35 |
| *RELN* | chr3 | 129923111 | 129923111 | 35 |
| *TRIO* | chr6 | 14733837 | 14733837 | 35 |
| *NTRK1* | chr1 | 94089606 | 94089606 | 34 |
| *MTOR* | chr1 | 213265857 | 213265857 | 34 |
| *RAI1* | chr16 | 17644443 | 17644443 | 34 |
| *RELN* | chr3 | 129793522 | 129793522 | 34 |
| *CD38* | chr5 | 14788455 | 14788455 | 34 |
| *TRIO* | chr6 | 14752487 | 14752487 | 34 |

### Cross-species comparison of highly constrained genes in humans and macaques

A number of genes implicated in human NDDs were characterised by high constraint scores in both humans and macaques, as evidenced by their RVIS scores more than two standard deviations from the mean (N = 5 genes with macaque: RVIS < -1.47, human: RVIS < -1.97) [**Supplementary File 6:** “RVIS scores human and macaque overlapping NDD genes”]. These include *TRIO, RAI1, MTOR, TSC2* and *RELN*. Two of these genes are implicated in NDDs, namely *RAI1* (Smith-Magenis Syndrome) and *TSC2* (Tuberous Sclerosis) Both Smith-Magenis syndrome and Tuberous Sclerosis are known to be associated with moderate to severe intellectual disability in humans and are also characterised by other medical and neuropsychiatric features. These genes and their mGAP annotated predicted pathogenic mutations are illustrated in **Supplementary File 7** “NDD_variants_top5_rvis_human_macaque”. Interestingly, a further four NDD genes were characterised by high constraint scores in humans but not in macaques. These include *CACNA1D*, *CNTNAP2*, *MBD5*, *AUTS2* and *NRXN1*. (**Supplementary File 8** - NDD_variants_top5_rvis_human_notmacaque).

## Discussion

In this study, we investigated the mutational burden in ASD and other neurodevelopmental genes among macaques in two fundamental ways. First, we determined whether genes known to be implicated in ASD and a number of related neurodevelopmental disorders in humans show evidence of evolutionary constraint against the accumulation of mutations. Second, we examined whether any of those macaque mutations that are predicted damaging in humans overlapped with known ASD and other NDD-implicated mutations in humans. Our analysis identified evidence of evolutionary constraint against the accumulation of mutations across all ASD and other NDD gene sets, indicating that, in macaques, as in humans, these genes have important functions. We also demonstrated that predicted damaging mutations were still quite common in all NDD genes, including those that overlap across all NDD disorders and are presumably the most important. Again, given their predicted damaging nature, it might be anticipated that these variants would have phenotypic consequences.

Warren et al. (2020) (30) examined sequence diversity in a sample of 853 macaques and identified thousands of missense variants across the genome impacting genes associated with human disease. Our results are largely consistent with their study, which analysed sequence diversity in this same dataset. Unlike our study, which set out to examine constraint across neurodevelopmental genes in different ways, theirs specifically examined the frequency of missense variants in these genes compared to all genes and found them to be depleted for missense variants. Nevertheless, our investigation did identify the fact that these NDD genes do harbour missense and other potentially damaging variants that could have phenotypic consequences, as discussed subsequently.

It is perhaps unsurprising that NDD genes show evolutionary constraint similar to humans. These genes clearly play a key role in core brain functions, as evidenced in our network analysis, and as such are presumably central to developmental brain processes. Given that macaques are among our most recent NHP ancestors, we might anticipate that genes that are important in humans would also be so in macaques. However, given that the neurodevelopmental disorders studied are all characterised by language abnormalities (including language pragmatics), which are evolutionarily advanced skills that characterise the very essence of humanness, there may also be a reason to have predicted that these genes would be less important (and hence less constrained) in macaques if their clinical impact is indeed at this higher cognitive level. Given our findings, the role of these genes in brain development is more likely to be pre-linguistic and hence at a far more fundamental level than the traits that characterise these disorders. Consequently, these genes may play a less important role in the ‘high-level’ manifestations of these disorders, which include aspects of social skills and language itself. What ultimately determines the clinical phenotype itself is likely to be downstream of the genes through their interaction with molecular neurobiology, neural circuits and connectivity through environment and other, as yet undetermined, factors.

We also demonstrated that, despite constraint, macaques harbour predicted damaging mutations in key NDD genes. These predictions are based on mGAP’s own annotation, which is derived from multiple sources of information after variants are lifted to the human genome, including established prediction tools (such as CADD and SIFT) and ClinVar annotations. These were supplemented with SnpEff annotation for predicted impact on protein-coding genes. Given this, in humans, these variants would be expected to have phenotype consequences, although both variable expressivity and variable penetrance will both confound the prediction of phenotypic consequences.

Unfortunately, phenotypes are unavailable for many of the macaques studied, so the impact of these mutations at a behavioural level is unknown. However, It remains possible that there are similar phenotypic consequences to humans among macaques at a neurofunctional, neurophysiological and/or ‘clinical’ level. This highlights the further insight that would be gained from collecting such data on NHPs, which could be relatively straightforward and largely non-invasive given the availability of such tools as eye tracking, EEG, neuroimaging and machine learning approaches to the collection and classification of behavioural observations.

Importantly, studies have already shown that macaques share remarkable similarities between human and macaque socio-cognitive systems (4,9,31). Neurobiological studies in humans and macaques have already shown evidence for evolutionarily conserved fronto-temporal networks that are particularly important for language that cannot be modelled in the same way using other experimental animals (32–37). Previous studies using eye-tracking methods showed that, similar to typically developing infants, macaques attend preferentially to social stimuli (7,8). Recent macaque genetic studies have also shown that macaques with single gene editing or naturally-occurring mutations present a combination of phenotypic behaviours including social and learning impairment as well as repetitive behaviours (18,20,38).

One further insight relates to the observation that a number of genes with strong evidence of constraint in humans were not similarly constrained in macaques. Some of these genes, such as *CACNA1D*, *CNTNAP2*, *MBD5*, *AUTS2* and *NRXN1* are key neurodevelopmental genes with high penetrance for clinical phenotypes. For example, copy number variation (CNV) in *NRXN1*, which encodes a synaptic scaffolding protein, has a penetrance of 6.4% for schizophrenia and 26% for ASD or ID (39). *MBD5*, a transcriptional regulator is implicated in ID, ASD, epilepsy and specific cognitive impairments (40). CACNA1, which encodes one of the L-type calcium channels is similarly implicated in ASD, ID and epilepsy. Crucially, these genes encode important aspects of brain structure and function, so their constraint against mutations in humans, and the phenotype consequence contingent on mutations occurring, is unsurprising. It is curious, therefore, that similar constraint is not seen in macaques. Gao et al. (2023) also identified a small number of variants with strong evidence of pathogenicity in humans that appeared to be well-tolerated in NHPs, proposing that interactions within the genomic neighbourhood may be relevant. Another possibility is that constraint is more fine-grained than at the gene level. For example, previous research has examined patterns of constraint among Pfam protein domains, showing tolerance to be consistent across specific domains in different genes, and that evolutionary conservation is correlated between entropy measured from Pfam and ExAC (41). Moreover, among some ID-associated genes, pathologic *de novo* mutations have been shown to cluster in specific protein domains (42). Examining patterns of constraint in different ways among primate datasets may therefore offer further insight into pathophysiology in human NDDs.

Human studies have identified many genes implicated in these NDDs, however, much is still not known about the neurophysiological, cognitive and other intermediate phenotypes. There are several limitations to human studies that may be overcome by shifting the focus to NHP-based research. One challenge has been the involvement of less able individuals with ID, ASD or epilepsy, who may be unable to follow instructions or comply with neuroimaging and electrophysiology, which are important tools for studying NDDs. It is important that such individuals are included in research, as they often represent the ‘purest’ forms of the disorder among whom there is wide consensus regarding diagnosis. However, recruiting such individuals and conducting research in an ethical manner has also been difficult, given the restrictions inherent in any research protocol.

New models are therefore needed. Up to now, NNDs have largely been studied in rodent models. However, rodents and humans have diverged significantly more than 70 million years ago, there are major differences in brain structure and function between the two species. This makes it difficult to recapitulate the research paradigms that capture clinical phenotypes for NNDs in rodents. For example, emotion recognition and processing of social information are important aspects of ASD and schizophrenia, but these are difficult to measure in rodents. In short, rodent models may be useful to capture cognitive intermediate phenotypes, however, they are almost certainly impossible to use to capture social-cognitive functions that require the primate-specific frontal structure through eye tracking, gaze and arousal in response to salient stimuli.

NHP models offer many advantages over rodent models for studying NDDs. Macaque is a particularly promising model for studying NDDs as they share many similarities with humans in terms of social behaviours, and they have a well-characterised genome. With the availability of a reference genome and the possibility of studying naturally occurring mutations in colonies, the challenge now is to develop research paradigms that capture the characteristics of neurodevelopmental disorders in both clinical terms and in relation to the neurobiology of cognition that are immediately downstream of these clinical manifestations. To this end, recently the macaque version of the Social Responsiveness Scales (mSRS) (14,15) has been developed. The mSRS is a screening questionnaire that can identify animals who demonstrate vulnerabilities indicative of ASD, based on the SRS which is widely used in clinical practice as a screening tool (43). The usefulness of the mSRS has been illustrated by several studies, for example, significant cerebrospinal fluid (CSF) arginine vasopressin (AVP) was reported in low-social macaques as seen in individuals with ASD (44) and the associations between social behaviours and whole genome sequence data in macaques (45). ‘Translating’ the human characteristics of disorders such as ASD into animal behaviour is not straightforward, however, and it is hoped that further work in this area will now thrive in anticipation of the major contribution of NHPs in biomedical science.

## Limitations

The unavailability of behavioural and other phenotypes in the macaques studied limits the conclusions that can be drawn regarding the possible consequences of the pathological mutations described. Furthermore, other measures of evolutionary constraint are available, although research supports consistency between these different metrics. There are also limitations with gene lists used for particular disorders, given that these lists change over time with the accumulation of new evidence. It might be argued that these lists are overly inclusive. To circumvent this, we conducted analyses for genes that overlapped between disorders, representing a more robust group of genes.

## Conclusions

The presence of pathological mutations in NDD genes among macaques, and the evidence of similar constraint in these genes to humans, provides a strong rationale for further modelling genotype-phenotype relationships in macaques. Given their ability to model social cognition and other higher cognitive functions, and their more recent evolutionary separation from humans, macaques are an ideal model for translational purposes to further our understanding of the neurobiological underpinnings of ASD and other NDDs. Macaques also offer the opportunity to explore the nature of this relationship through mediating neuropsychological and neurofunctional modelling. This highlights the importance of this species as a prominent animal model to understand the neurobiological underpinning of ASD.

## Abbreviations

ASD: Autism spectrum disorder
NDD: Neurodevelopmental disorder
ID: intellectual disability
RVIS: Residual Variation Intolerance Scores
GSEA: Gene Set Enrichment Analysis
NHP: nonhuman primate
mGAP: macaque Genotype and Phenotype
SRS: Social Responsiveness Scale

## Acknowledgements

mGAP is supported by NIH (R24OD021324).

## Author contributions

YK and MWS conceived the study and undertook data analysis. YK, MU, CM, JV, SW and MWS, all undertook interpretation of data and preparation of the manuscript.

## Declarations

## Consent for publication

Not applicable

