## Supplemental materials for "Evolutionary constraint genes implicated in autism spectrum disorder across 2,054 nonhuman primate genomes"

Supplementary Figure 1: Correlation of human and macaque RVIS scores.
Supplementary Table 1. Summary statistics for all macaque variants before and after QC
Supplementary Table 2: RVIS for different clinical gene set
Supplementary Table 3: Gene over-representation analysis top 2% constrained genes
Supplementary Table 4: GSEA by diagnosis

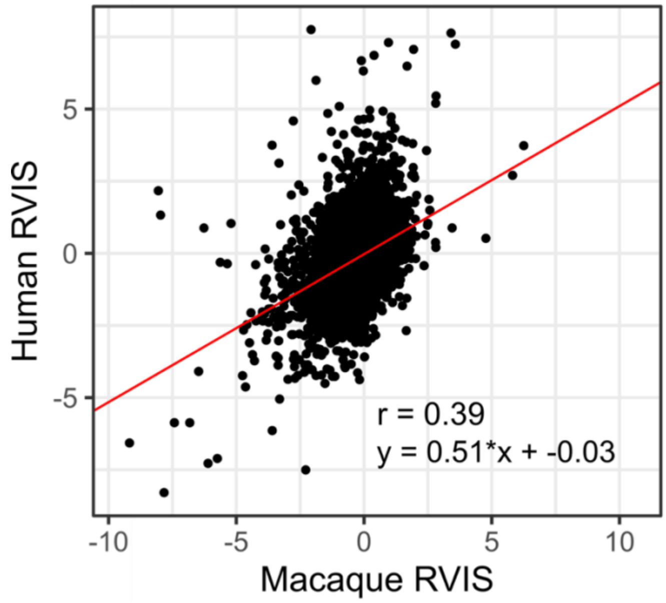


**Supplementary Figure 1.** Correlation of human and macaque RVIS scores. The correlation was calculated using the macaque and human data (25) across 11898 overlapped genes between macaque and human data after removing the outliers (RVIS > 11). The 95% confidence area ([0.38 - 0.41]) was calculated with a fitted liner regression (red line: Pearson's product-moment correlation = 0.39, $df$ = 11,896, $P < 2.2e^{-16}$).

**Supplementary Table 1. Summary statistics for all macaque variants before and after QC**

| | Pre QC | Post QC |
|---|---|---|
| Total variants | 40,391,636 | 37,503,663 |
| Total SNPs | 34,741,086 | 32,507,418 |
| Total Indels | 5,650,550 | 4,996,245 |
| Frameshift mutations | 28,102 | 21,584 |
| Missense mutations | 418,065 | 370,683 |
| Stop gained mutations | 12,757 | 10,035 |

**Supplementary Table 2: RVIS for different clinical gene set**

| | Mean RVIS (SD), number of genes | Mean RVIS (SD), background no. genes | t | p | β (std. err) | Odds Ratio |
|---|---|---|---|---|---|---|
| Overlapping genes across all four NDDs | -0.52 (0.75) N=101 | 0 (0.68) N = 18,065 | 6.93 | $4.0 \times 10^{-10}$ | -0.53 (0.07) | 0.59 |
| ASD-DISGENET | -0.32 (0.88) N = 793 | 0.01 (0.67) N = 17,373 | 10.44 | $< 2.2 \times 10^{-16}$ | -0.49 (0.04) | 0.61 |
| ASD-SFARI | -0.44 (1.10) N = 729 | 0.02 (0.66) N = 17,437 | 11.13 | $< 2.2 \times 10^{-16}$ | -0.63 (0.04) | 0.53 |
| Epilepsy | -0.33 (0.79) N = 872 | 0.01 (0.68) N = 17,294 | 12.53 | $<2.2 \times 10^{-16}$ | -0.51 (0.04) | 0.60 |
| Schizophrenia | -0.25 (0.85) N = 2,021 | 0.03 (0.66) N = 16,145 | 14.40 | $<2.2 \times 10^{-16}<$ | -0.50 (0.03) | 0.61 |
| ID | -0.30 ( 0.84) N = 1,573 | 0.03 (0.66) N = 16,593 | 14.78 | $<2.2 \times 10^{-16}$ | -0.54 (0.03) | 0.58 |

**Supplementary Table 3: Gene over-representation analysis among top 2% constrained genes (N=369)**

| | Number of genes | Number overlapping | P-value | FDR |
|---|---|---|---|---|
| ID | 1570 | 89 | $1.7 \times 10^{-19}$ | $4.3 \times 10^{-19}$ |
| ASD | 793 | 50 | $9.6 \times 10^{-13}$ | $1.2 \times 10^{-12}$ |
| ASD-SFARI | 729 | 65 | $2.8 \times 10^{-24}$ | $1.4 \times 10^{-23}$ |
| Epilepsy | 871 | 51 | $8.9 \times 10^{-12}$ | $8.9 \times 10^{-12}$ |
| Schizophrenia | 2018 | 96 | $5.9 \times 10^{-16}$ | $9.8 \times 10^{-16}$ |

**Supplementary Table 4: GSEA by diagnosis**

| | Number of genes | P-value | FDR | ES |
|---|---|---|---|---|
| ID | 1570 | $2.1 \times 10^{-47}$ | $1.0 \times 10^{-46}$ | 0.19 |
| ASD | 793 | $2.7 \times 10^{-25}$ | $2.7 \times 10^{-25}$ | 0.19 |
| ASD-SFARI | 729 | $1.2 \times 10^{-33}$ | $2.0 \times 10^{-33}$ | 0.23 |
| Epilepsy | 871 | $7.0 \times 10^{-32}$ | $8.8 \times 10^{-32}$ | 0.21 |
| Schizophrenia | 2018 | $2.6 \times 10^{-44}$ | $6.5 \times 10^{-44}$ | 0.17 |
